# Save GPU RAM Usage in Convolutional Layers to Load Larger Histopathological Images

**DOI:** 10.1101/2023.09.19.558533

**Authors:** Takumi Ando

## Abstract

Image recognition models have evolved tremendously. Despite the progress for general images, histopathological images are not easy targets. One of the reasons is that histopathological images can be 100000-200000px in height and width which are often too large for a deep neural network model to handle directly because the RAM of the GPU is limited. Mitigating the obstacle is expected to be a progress in the histopathological image analysis. In this study, we save the RAM consumption in a convolutional layer by allocating only the required data to GPU only when needed and by dividing the calculation into per channel. This RAM Saving Convolutional layer (RSConv) can load larger images than a normal convolutional layer. The code is available at https://github.com/tand826/RAMSavingConv2d.

## 1 Introduction

Deep neural networks have proved their possibility in histopathological image analysis. The technical advancements can greatly help a daily routine diagnosis because of the labor-intensive nature of pathologists works through the microscope such as counting the number of cells in a specific region. The possibility is not limited to those works and recent research is trying to show the power even for a diagnosis of benign or tumor.

Digital histopathological images (Whole Slide Images, WSIs) are often huge because they need to represent the figures of cells. For example, when a pixel represents 0.5 microns, a 1cm tissue can be a 20000x20000 pixels image which represents a cell with around 20x20 pixels. This is often large for models to load on GPU compared to the size of general images. Previous studies have managed to load WSIs by patching them into small pieces. Methods of multiple instance learning (MIL) based on patching are now the thriving approach to get the feature of a WSI from a set of features of patches.

In models for general images, a traditional model often starts with a convolutional layer. It can reduce the input size to almost half in width and in height and the following layers are required to handle smaller data. The success of these models does not apply directly to WSIs because of the limit of the RAM. The optimization of the convolutional layer in the first layer is important for huge images to use popular models of general images in the same way.

In this study, we reduce the RAM consumption in a convolutional layer for 2D images at the expense of time to enable models to handle larger images by temporarily allocating the data on CPU or disk unless needed for calculations and by using minimal data on GPU for the convolution operation. Contributions are as follows. (1) We propose RAM Saving Convolutional Layer (RSConv) to reduce the RAM usage of a convolutional layer. (2) We evaluated the maximum loadable size, the RAM efficiency, and the improvement of the low time efficiency of RSConv by optimizing the calculation.

## 2 Related Works

Traditionally, WSIs are split into a number of small patches to avoid out-of-memory error and apply the paradigm of deep learning models for general images to histopathological images [6, 3]. For a WSI-level result, patch-level results are used to be aggregated by non-trainable operations. As opposed to per-patch models, MIL models also have been the subject of research. While Attention-Based MIL [7] proposed a fully trainable MIL model for histopathological images with a global attention pooling layer, [1] used MIL to get features of WSIs followed by aggregation with RNN. Recently, approaches focusing on the multiresolution structure of WSIs have been proposed, such as a model to extract features from multiple resolutions by per-resolution models pre-trained by self-supervised learning [2]. These patch-based approaches remain to be promising, although they have the disadvantage of not being able to use models for general images as is.

To handle larger images, [13] tried to train a huge image with its size kept as large as possible by disassembling ResNet [5] to blocks and training the model per block. They proposed a way to use general models for WSIs with Attention-based MIL as an extra module to optimize the blocks. [9] proposed a streaming approach, which is to disassemble a model into blocks with each intermediate feature deleted just after its computation and recalculate them when needed for backpropagation at the expense of time. In addition to the streaming approach, [4] added packing the input images into a smaller size rectangle by concatenating minimal bounding rectangles of tissues and minimizing the background area.

Some experiments tried to reduce GPU RAM usage for CNN-based models by focusing on the whole architecture [12, 8]. CPU offload strategy of deepspeed [11, 10] stores model parameters to CPU RAM or disk to save GPU RAM usage to train on GPUs with limited RAMs.

## 3 Methods

### 3.1 Forward Operation

RSConv keeps values to CPU or Disk in general and loads required values only when needed, whereas a normal convolutional layer keeps all the values on GPU. After the GPU receives the input value, RSConv moves it to CPU and allocates the memory for the output value on the GPU followed by the calculation of the output per input channel. In total, the peak RAM consumption on GPU is the sum of 1/N of the input value and output value for N channel images. After the calculation, the input value is kept on CPU for backward calculation.

### 3.2 Backward Operation

A convolutional layer calculates 3 parameters in backward operation: the weight gradient, the bias gradient, and the input gradient.

#### Weight Gradient

RSConv receives the gradient of the output feature map from the following layer and copies the input value kept in forward operation. With these 2 parameters, split the input values and the output gradient into per channel values, RSConv calculates the gradient weight with the peak RAM consumption limited the output gradient and 1/N of input value for N channel images.

#### Bias Gradient

The required parameter is only the output gradient. This consumes only the output gradient which requires less than the operation for the weight gradient.

#### Input Gradient

The input gradient requires the weight values and the output gradient. RSConv calculates the input gradient per output channel to limit the peak RAM consumption to 1/M of the output gradient and the input gradient where the output gradient has M channels of features. In this operation, the RAM for the result of the input gradient is not required to be allocated on CPU as the result of the calculation is on GPU when computed on GPU for faster operation. For acceleration, if the layer is the first layer of the model, RSConv skips the calculation of the input gradient because the value is not usually required to train the model. In addition, the number of channels in one calculation can be 2 or more until the consumption reaches the limit as the output feature size is often smaller than the input shape.

## 4 Materials

We decided to test RSConv with a 22182x22182 image concerning The Genome Cancer Atlas (TCGA) dataset.

The total number of WSIs of TCGA is 29888 including the diagnostic and the tissue slides and excluding the ones with unreadable properties. The median size of the WSIs is 77111px in width and 34294px in height. As a 16x16 feature of 20x WSIs fits the size of a single cell as in [2], we choose 20x magnification. By reducing the blank area by packing as [4], the size is 23937px in width and 20556px in height. 22182x22182 is the nearest integer size to have the same area as the median image above.

## 5 Experiments and Results

### 5.1 Experimental Setups

All the experiments are employed on a GPU workstation with RTX3090 (24GB GPU RAM) and 64GB CPU RAM and Python 3.9.9 environment built with CUDA 11.7 and PyTorch 2.0.1.

Convolutional layers are set to output a feature of 16 channels with 7 pixels width and height of kernel strides with 2 pixels. Images to feed to the layers are in float16 of RGB color space and padded 3 pixels on calculation.

### 5.2 RAM Efficiency

For an input image with 3 channels of 22182x22182, the output feature is 16 channels of 11091x11091. While the forward calculation of a normal convolutional layer requires the sum of the input and output sizes, RSConv requires those of a single channel of the input and the output. The backward calculation of a normal convolutional layer requires the size of the input, the input gradient, and the output gradient. This is larger than the peak RAM consumption of RSConv which is the sum of a single channel of input and output gradient or the sum of a single channel of output gradient and the input gradient. Overall, the maximum RAM consumption of RSConv is smaller than a normal convolutional layer as shown in Fig 3. In addition, we tested ResNet18 with its convolutional layers changed with RSConvs, and Fig 4 shows the result of RAM usage reduction.

**Fig. 1.**
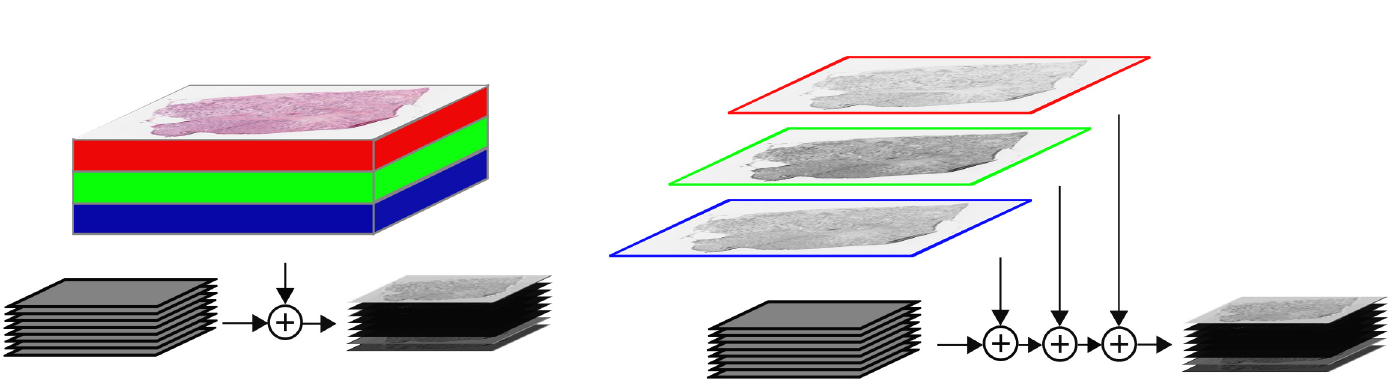
Convolutional layers for a WSI of RGB 3 channels to get a feature of 8 channels. Both add the output features to the allocated blank tensor. **Left**. The normal convolutional layer outputs the feature in one operation. **Right**. RSConv computes the features per channel and adds them one by one.

**Fig. 2.**
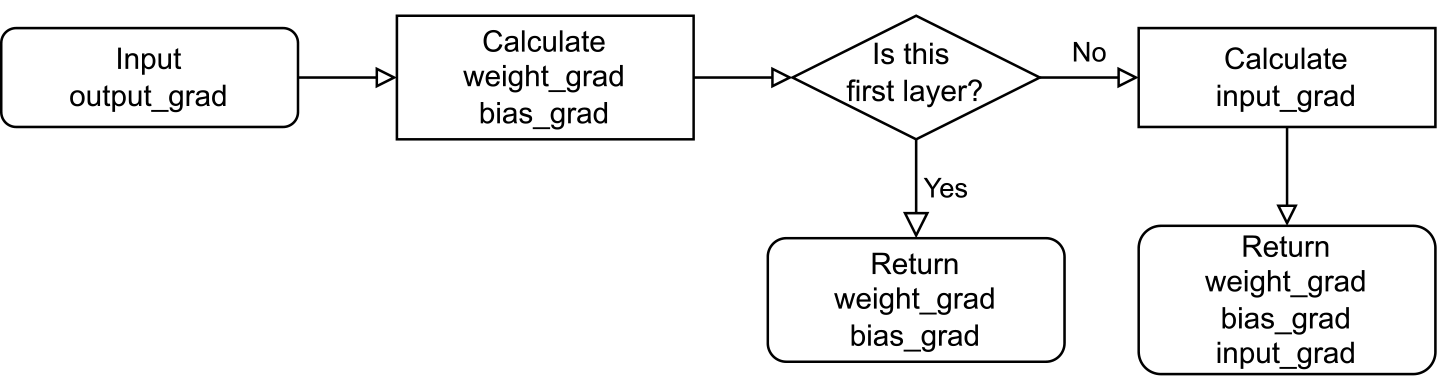
The general flow of the backward calculations of RSConv.

**Fig. 3.**
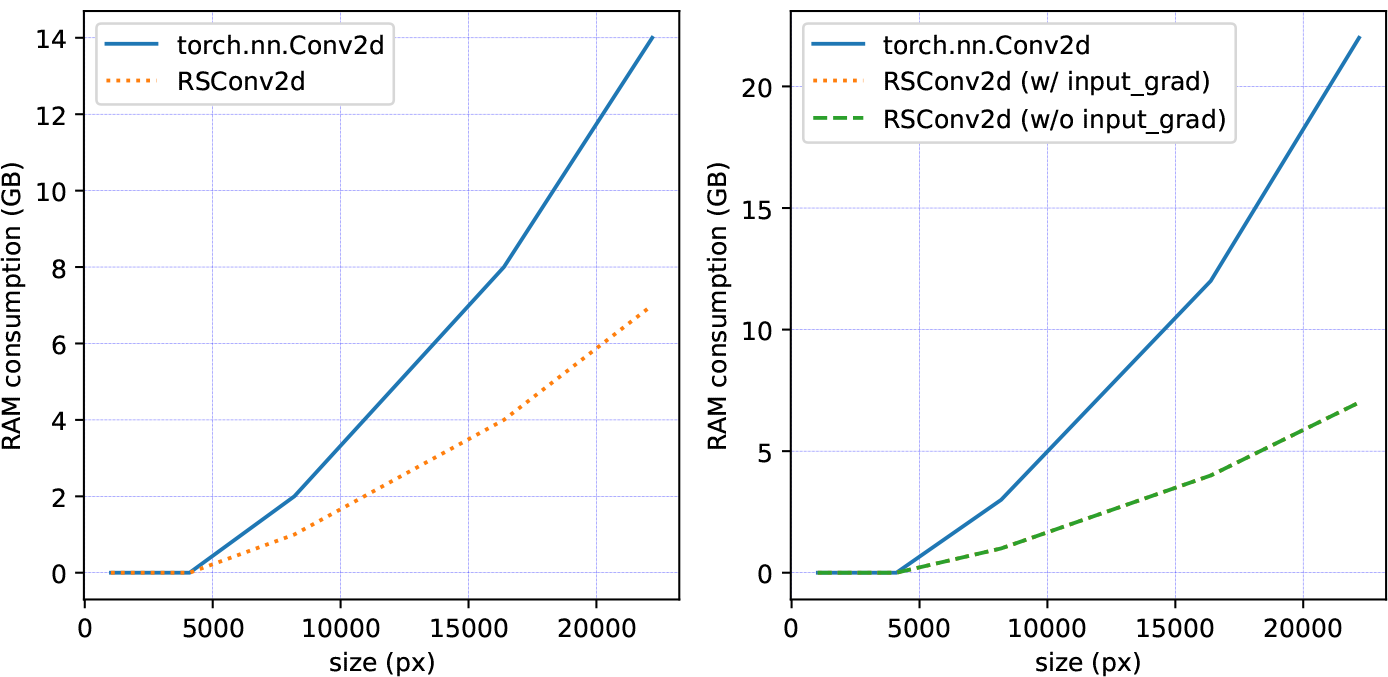
The peak RAM consumptions of convolutional layers (GB). **Left**. Forward operation. **Right**. Forward operation and backward operation with and without the calculation of input gradient.

**Fig. 4.**
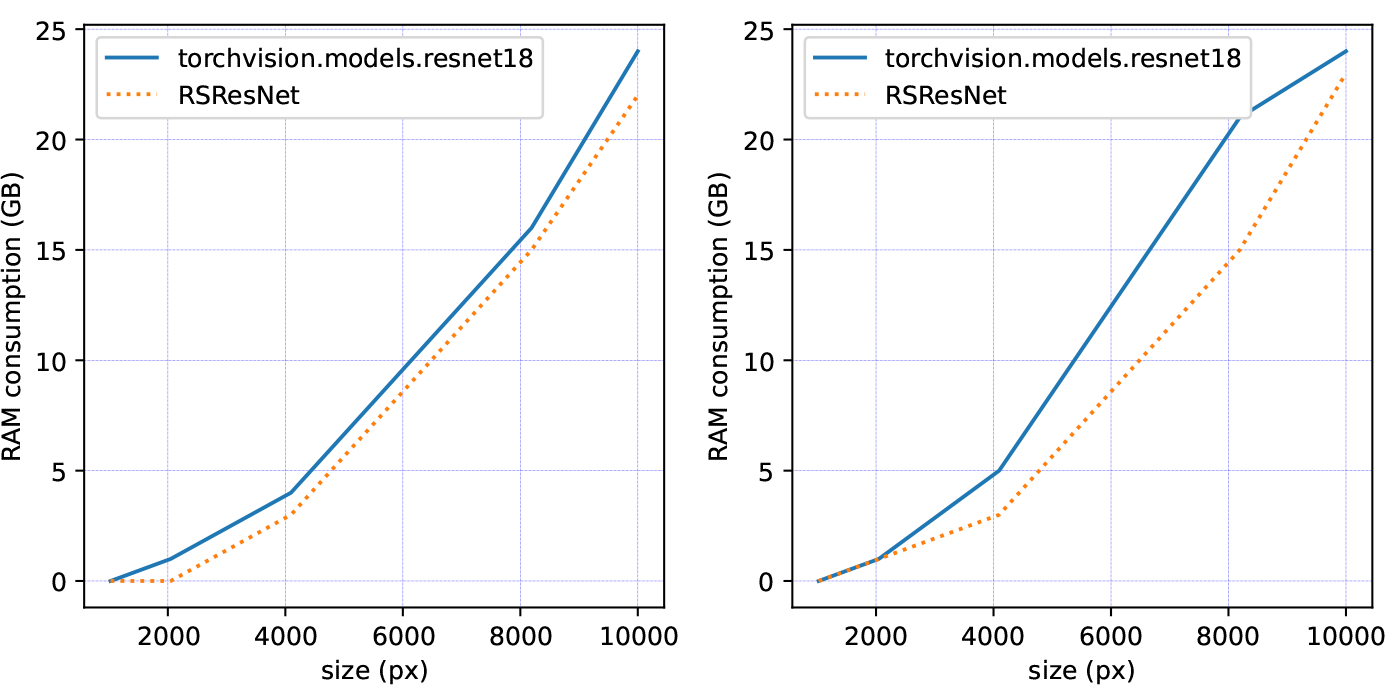
The peak RAM consumptions of ResNet18 with and without RSConv (GB). **Left**. Forward operation. **Right**. Forward operation followed by backward operation with the calculation of input gradient of the first layer skipped in ResNet with RSConv.

### 5.3 Time Efficiency

RSConv can skip the calculation of the input gradient if the convolutional layer is the first layer of a model to reduce the processing time. This is not available when used in the middle of a model. Also, the saved time is wasted by the copying time between the devices and the calculations repeated the number of the channels of output. The actual times are in Table 1.

**Table 1.** The processing times of convolutional layers (milliseconds).

| Layers | Forward | Forward<br>+Backward(w/ input grad) | Forward<br>+Backward(w/o input grad) |
| --- | --- | --- | --- |
| torch.nn.Conv2d | 0.21 | 185.756 | N/A |
| RSConv | 2736.65 | 9339.78 | 5309.08 |

### 5.4 Size Efficiency

We observed the maximum loadable size of convolutional layers. RSConv could handle the largest image when operating only the forward calculation and the forward calculation followed by the backward calculation skipping the input gradient, although a normal PyTorch convolutional layer constantly handles an image with 23000px in width and height as in Table 2.

**Table 2.** The maximum size of images for convolutional layers when used as a single layer (pixels).

| Layers | Forward | Forward<br>+Backward(w/ input grad) | Forward<br>+Backward(w/o input grad) |
| --- | --- | --- | --- |
| torch.nn.Conv2d | 23000 | 23000 | N/A |
| RSConv | 34500 | 26500 | 34500 |

## 6 Conclusion

We could save the RAM consumption in a convolutional layer to load huge images with the speed loss kept minimal, especially in the first layer of a model. While the operation is required once in a normal convolutional layer for RGB input image, RSConv repeats it 3 times. In the same way, the backward operation repeats the number of channels of the output feature. This results in a lower time efficiency in backward operation. Taking advantage of skipping the input gradient calculation in the first layer, it is desirable to use RSConv only in the first layer. The optimization after the 2nd layer or later remains an issue.

## Acknowledgements

The results shown here are in part based upon data generated by the TCGA Research Network: https://www.cancer.gov/tcga. This research is funded by PSP Corporation, SoftBank Corp., Nikon Corporation, Sakura Finetek Japan Co., Lid., Roche Diagnostics K.K., Sysmex Corporation, Preferred Networks, Inc., NEC Corporation, SEIKOTEC CO., LTD., CGI, K K, Hologic, Inc..

